# A Compartmentalized Model to Directional Sensing: How can an Amoeboid Cell Unify Pointwise External Signals as an Integrated Entity?

**DOI:** 10.1101/2022.10.24.513450

**Authors:** Zahra Eidi, Mehdi Sadeghi

## Abstract

After being exposed to an external chemical attractant, several internal cellular downstream signal transduction pathways control the chemotactic machinery of eukaryotes. The spatial activation of these pathways ultimately leads to some sort of symmetry breaking around the cell periphery in the form of redistribution of several biochemicals, *e.g*., polymerized actin at one side of the cell for propulsion and the assembly of myosin II at nearly opposite side for retraction. In this study, we revisit the modeling of this process, which is called directional sensing, with a compartment-based design. In our model, we consider a network of excitable elements around cell circumference which are stimulated occasionally with local colored noise. The exciting elements are capable of sharing information with their close neighbors. We show that the dynamic can distinguish a temporary but enough long-lasting direction, statistically, pointing towards the gradient of external stimulants that can be deemed as the preferred orientation of the cell periphery during the directional sensing process in eukaryotes.

## Introduction

Directional sensing is an influential feature in eukaryotic chemotaxis. The ability possesses the asymmetric recruitment of intracellular proteins to the cell membrane when a cell is exposed to a gradient of its chemoattractant. This incorporation requires the coordination of complex mechanisms on the microscopic level, including transducing information from the external ambiance to the interior of the cell and translating it to a migratory response via a molecular signaling pathway based on the environmental cues [1]. The amoeba *Dictyostelium discoideum* is a well-established model organism for studying amoeboid locomotion and chemotaxis in eukaryotes. The cell extends successive pseudopodia, a temporary actin-based protrusion of its surface, in order to propel on the surface following the signaling messenger cyclic Adenosine Mono Phosphate (cAMP) gradient [2]. The cells are pretty keen to sense and respond to the chemical stimulants such that they are capable of detecting a one percent difference in concentration of the chemoattractant from one side of their circumference to the other, over a wide range of steepness [3]. But, how the cell can unify the information of detected local signals around its body surface to the global response as an integrated entity?

Based on experiments large interconnected or/and parallel [4] networks of signal transduction events triggered within seconds of chemoattractant exposure [5]. During this time, the starved cells demonstrate some special feature of symmetry breaking within the cell in which PI3K molecules are localized in leading edge, where the gradiant of stimulant concentration is higher, and PTEN molecules at some point roughly across from the leading front of the cell even when they are immobilized [6,7]. Experiments show that cells essentially respond to external chemical stimulant cAMP in a wide range of its concentration gradient and this response occurs in variant background cAMP concentrations (adaptation) [3,8]. Besides, the induced front-back asymmetry inside the cell is steeper than the external gradient of chemoattractant concentration (amplification) [6]. Moreover, whenever the gradient direction is reversed, the position of front and back are exchanged in such a way that the cell gets aligned with the new direction (re-orientation) [9].

Based on the “classic” view of directional amoeboid locomotion, signal transduction is merely a bridge-like process upon which the external information of receptor occupancy is translated to cytoskeletal motility responses. However, recent observations imply more interconnected dependency between these processes [10]. Demonstration of actin waves [11,12] and the spinning waves of PIP3 and PTEN [13] are pieces of evidence, among others [10], that suggest more elucidate models are needed in this area. Assembling these components to form a clear picture of the whole process still needs formal approaches to make the research in this field, inevitably, productive and sensible.

Local Excitatory and slower, Global Inhibitory process (abbreviated as LEGI) was one of the earliest and of course most fitted models for the spatiotemporal responses of immobilized cells. The model extended to explain gradient sensing in migrating cells, the ability of a living organism to produce adapted and amplified responses, traced back to a fast and local excitatory reply that accompanies a slow and inhibitory signal [14–16]. In more recent versions of LEGI-based models, parallel mechanisms are introduced to regulate the accumulation of PI3K and PTEN proteins in the front and back of the cell [17]. The appearance of features like actin waves and PIP3 waves displays properties of excitable biochemical systems. The properties mostly exhibit all-or-nothing responses to suprathreshold stimuli and a refractory period. In order to describe the features, signal transduction excitable network (STEN) model was proposed to explain transient shifts between the rear and leading edges of the plasma membrane [10,18]. This in turn can coordinate cytoskeletal excitable network (CEN) to generate pseudopodium that mediates cell movement [19]. It has been shown that incorporating STEN model with LEGI module in the so-called biased excitable network (LEGI-BEN) can justify most of the temporal and spatial responses to chemoattractant [20]. Research in this area has had a paradigm shift, substituting the previous mechanism-based modeling with an advanced mathematical description of amoeboid motility, envisioning a more systematic, data-driven approach [21].

In this study, aiming to incorporate variant aspects of self-organizing activity in the signal transduction process, we consider that the signaling transduction system is near the threshold for activation in an excitable circumferential network. The compartments of this network are being reduced to two main players each of which shares information with its corresponding agent in an adjacent neighbor. However, the coupling net impact of these two reduced players, the activator and the inhibitor are opposite: While the activator variable is inclined to congregate, the inhibitor one tends to diffuse. Besides, we assume that the driving force to excite the network is provided by an additive local colored noise. We see that the setup can result in the accumulation of activators and inhibitors in two distinct and distant points of cell circumference. Remarkably, the activator accumulation point is often located at locally higher concentrations of external stimulants in an inhomogeneous medium, suggesting that the point at which the activator is highest picks out the leading edge of the cell in the medium.

## Materials and methods

The living entities are not purpose-built artifacts, engineered to consistently perform an exact action in response to an incoming signal; rather a cell is more akin to a soup of ingredients colliding and aggregating within cells. Local events in this animate soap should result in global actions, thus the key to deciphering the behavior of a living system is to understand how the collective effects of these local events can give rise to the response of an inanimate artifact with global rules.

We are about to model the global response of a cell in response to environmental chemical stimulants, Fig.1(a). To capture the compartmentalized nature of a living cell, we assume that the circumference of our artificial circular cell is comprised of a set of coupled locally excitable compartments. The size of these compartments is such that a well-stirred environment for the chemical reactions occurring within them is available. Assuming that an excitable element consisting of an activating variable *A* and an inhibiting one *I* are representative features of the resulting local cascade of chemical events within each element, we construct the model as a spatially organized ensemble of excitable elements which are located on a ring lattice with *N* elements, Fig.1(c). We assume that the below spatially discretized dynamic governs the evolution of the concentration of *A* and *I* in each element:

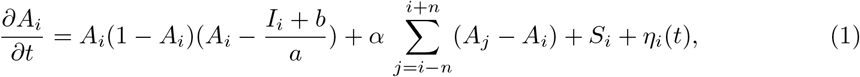

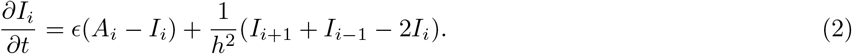

where *h* is the grid spacing. The index of the dynamical variables *A_i_* and *I_i_* with *i* = 1,…, *N* determines the element location in the lattice. Here, the excitable nature of each element on the lattice is described by a modified version of *Barkley* model [22]. We assume that within the interval [0.*δ*) activator variable *A* vanishes,i.e. in every cell whenever *A_i_* < *δ*, its value in the next update will be equal to zero. The values of the variables are updated using a discrete-time derivative *Euler* method. All the elements are identical and each of them is coupled with the other elements via the second terms of Eq.1 and Eq.2. The integer *n* defines the nonlocality of the coupling, which we call it the interaction interval. The coupling dynamic of *A_i_* with its neighboring corresponding values, the second term of Eq.1, implies that the concentration of *A_i_* is going to be amplified provided that its current value is higher than that of its neighbors. On the other hand, the second term of Eq.2 is nothing but a diffusion-like dynamic that tends to subside *I_i_* concentration in the next step, if its current value is higher than the local concentration of it in its adjacent vicinity. We perform our simulations on a lattice with N compartments with periodic boundary conditions.

**Fig 1.**
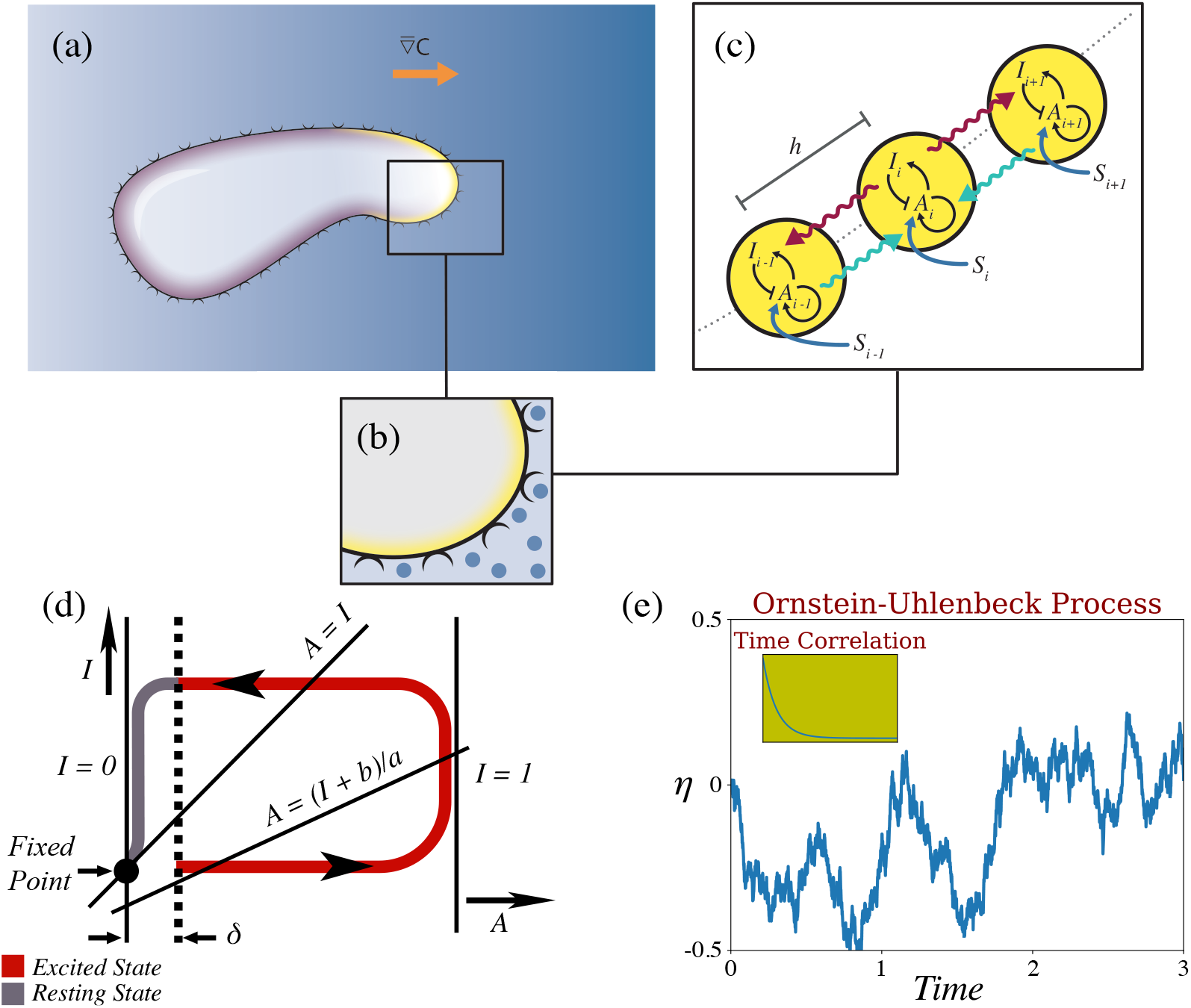
(Color online) (a) The illustration of a cell moving to the right of an inhomogeneous medium in response to a chemoattractant gradient which is indicated by the blue-scaled background. (b) The binding of chemoattractants, depicted as blue points, to circumferential receptors triggers internal signaling pathways inside the cell. (c) Our model assumes the peripheral area of a cell is composed of *N* coupled locally excitable compartments, consisting of an activating variable *A* and an inhibiting one *I*. In our simulations *N* = 120. The conducted signal in each compartment, *e.g., S_i_*, stimulates these excitable systems. (d) Schematic illustration of local dynamics in each excitable compartment. Corresponding dynamics of the excitable system in *AI*-plane is depicted by its nullclines. *A* has three separated nullclines: *A* = 0, *A* = 1 and *A* = (*I* + *b*)/*a*, while the line *A* = *I* corresponds to *I* nullcline. The dotted line indicates the boundary layer *δ* which all the initial conditions within it subside to the fixed point. The long itinerary in the *AI*-plane starts once the value of *A* exceeds the amount of *δ*, adapted from [22]. (e) Depiction of a typical realization an Ornstein-Uhlenbeck process with relation 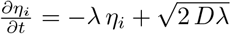 in which *D* = 0.5 and *λ*^−1^ = 1. The corresponding time correlation function of the process is shown in the inset.

The parameter *ϵ* indicates the excitability of the system which is the time-scale difference between the fast excitatory variable *A* and the slow refractory variable I. The parameters *a* and *b* are the controlling parameters of the model. This dynamic is characterized by a stable but excitable fixed point, see Fig.1(d) for a schematic illustration. Small perturbations from the rest state decay and return to the fixed point immediately. However, perturbations that exceed the boundary layer of size *δ* increase and settle downs on the fixed point only after the system has performed a large excursion in the *IA*-phase plane. Then, the system relaxes at the fixed point for the next incoming perturbation. The initial condition for the dynamic of *A_i_*s and *I_i_*s in the simulations are random values of the variables with a uniform distribution within the intervals: 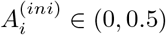, 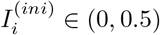.

In the full signaling circuit, when exposed to extracellular chemoattractants, the signals are detected by receptors on cell membrane, Fig.1(b). The resulting signal *S_i_* is funneled through several signal transduction modules, ultimately leading to higher concentration of activating agents at the front of cell where the amount of *S_i_* is higher, Fig.1(c). Throughout our simulations, We assume that *S_i_* is proportional to the probability of occupying the *i*-th compartment which is equal to 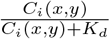. Here, *C_i_*(*x,y*) is concentration of external stimulant at the position of *i*-th compartment in *xy*-plane. By definition, for a concentration of chemoattractant equal to *K_d_*, half of the receptors of the cell are occupied [23]. However, in our simulations, *K_d_* is a fixed parameter equal to 0.5.

In Eq.1, the function *η_i_*(*t*) is the affected noise on the activator variable *A* on the *i–th* element. We assume that *η_i_* function is driven by a colored noise via an *Ornstein–Uhlenbeck process* in each compartment independently. The relation 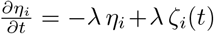 dominates the evolution of the random varible. Here, *ζ_i_*(*t*) is a Gaussian white noise of mean zero and correlation function < *ζ_i_*(*t*)*ζ_i_*(*s*) >= 2*D δ*(*t* – *s*). Fig.1(e) provides an illustration of a typical realization of this ‘mean reverting’ dynamic. The corresponding variance of this stochastic process and its correlation time are *D λ* and *λ*^-1^, respectively. Note that *λ* has units of frequency, and gives a measure of the cutoff frequency of the fluctuations of the noise; thus *λ*^-1^ gives a measure of its autocorrelation time.

To perform integration of η¿ function, in line with the integration of the deterministic part, we implement the *Euler–Maruyama* technique with a time step Δ*t*,

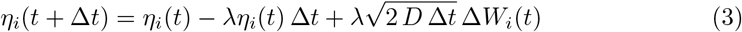

where the Δ*W_i_*(*t*)’s are independent Gaussian random numbers with unit variance [24].

## Results

We assume that a homogeneous medium is an environment within which the concentration of chemoattractant is uniformly distributed and the uniform value is constant, either zero or non-zero. Fig.2 illustrates the evolution of activator *A* and inhibitor *I* for all the elements of a cell which is located in a homogeneous medium, during the process of directional sensing on a single step. In these heat maps, vertical and horizontal axes represent time and indices of the elements, respectively.

**Fig 2.**
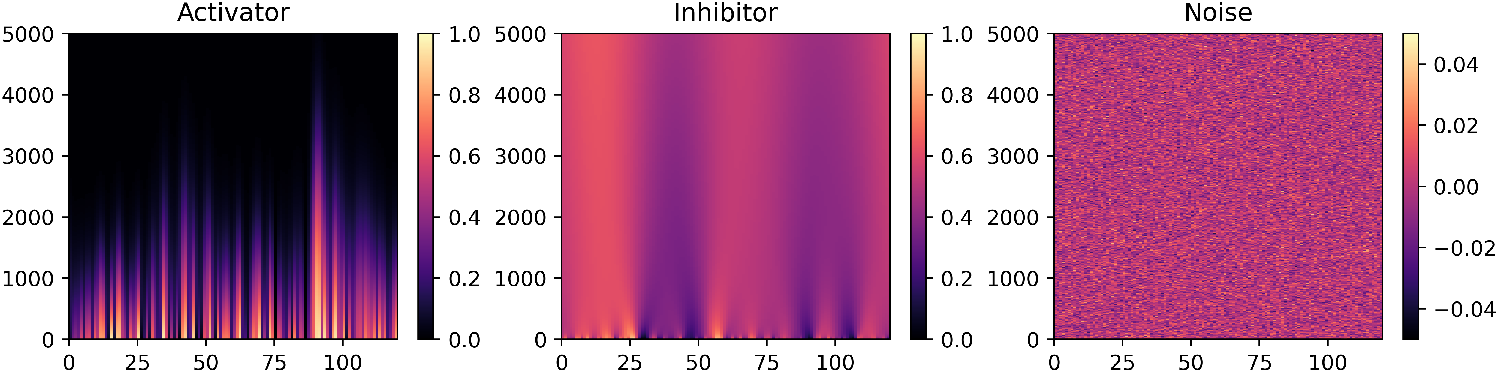
Heat map showing the concentration of (Left panel) activator concentration *A*, (Middle panel) inhibitor concentration *I*, and (Right panel) amount of noise *η* in each point of circumference during a single step(including 3987 iterations) in a homogeneous field. Note that both index 0 and index 120 address the same compartment and point to the right direction of horizon. Here, (a) the compartment with index 90 in which *A* outlasts more than other compartments, introduce the so-called “front” of cell and (b) the compartment with index 13 in which *I* outlasts more than other compartments, introduce the so-called “back” of cell.

### Homogeneous Medium

The period of this simulation has not been restricted to a special time interval. Rather, starting from a reliable random state, the simulation has been performed untill the value of *A* vanishes in each and every compartment. For example in Fig.2, the cell performed 3987 time steps, iteratively, to find its preferred direction at the end of the period.

Here, in a homogeneous medium, the noise is represented by *η* function. From the heat map of *η* in Fig.2, it is evident that at the beginning of simulation the function takes random values *η*_0_(*i*) for every compartment. In some elements, the initial value is enough for passing them through the *δ* barrier and entering in the excited phase. However, there are elements that have to return to their fixed point as the amount of their corresponding *η* is not sufficient to start an excursion in the *AI*-plane. It is worth mentioning that obviously the additive noises are not positive in all elements. The third panel of Fig.2 depicts the corresponding heat map of *η* function. It is seen that with a small perturbation coming from this noise, the internal dynamic in each element is triggered. That is why they call it noise-driven excited state dynamic. The coupling term in Eq.1, accelerates the process of activator aggregation in each step. We assume that the compartment in which A lasts longer than others specifies the so-called *front* of cell. In the corresponding heat map of *A* in Fig.2, one can follow the process of growing *A* variable in each compartment. The highest outlasting *A* in this heat map occurs at the element with index *j* = 90 which lasts for 3987 iterations. Evolution of *A*_90_(*t*), *I*_90_(*t*) and *η*_90_(*t*) is depicted in the first column of Fig.3. The middle panel of Fig.2 illustrates the evolution of *I* function on all the elements around the cell during the course of time. Evidently, activator and inhibitor concentrations evolve in nearly opposite direction, *i.e*. wherever *A* value is higher, the corresponding *I* value is lower, and *vice versa*. We define the *rear* side of a cell by the index at which the amount of *I* vanishes later, *e.g*., for the simulation of Fig.2, the corresponding element is locating at *j* = 13 which is seating at an acceptable distance from the front of cell located at *j* = 90, The angular separation of front and back in this circular cell is nearly 2.25 radians.

**Fig 3.**
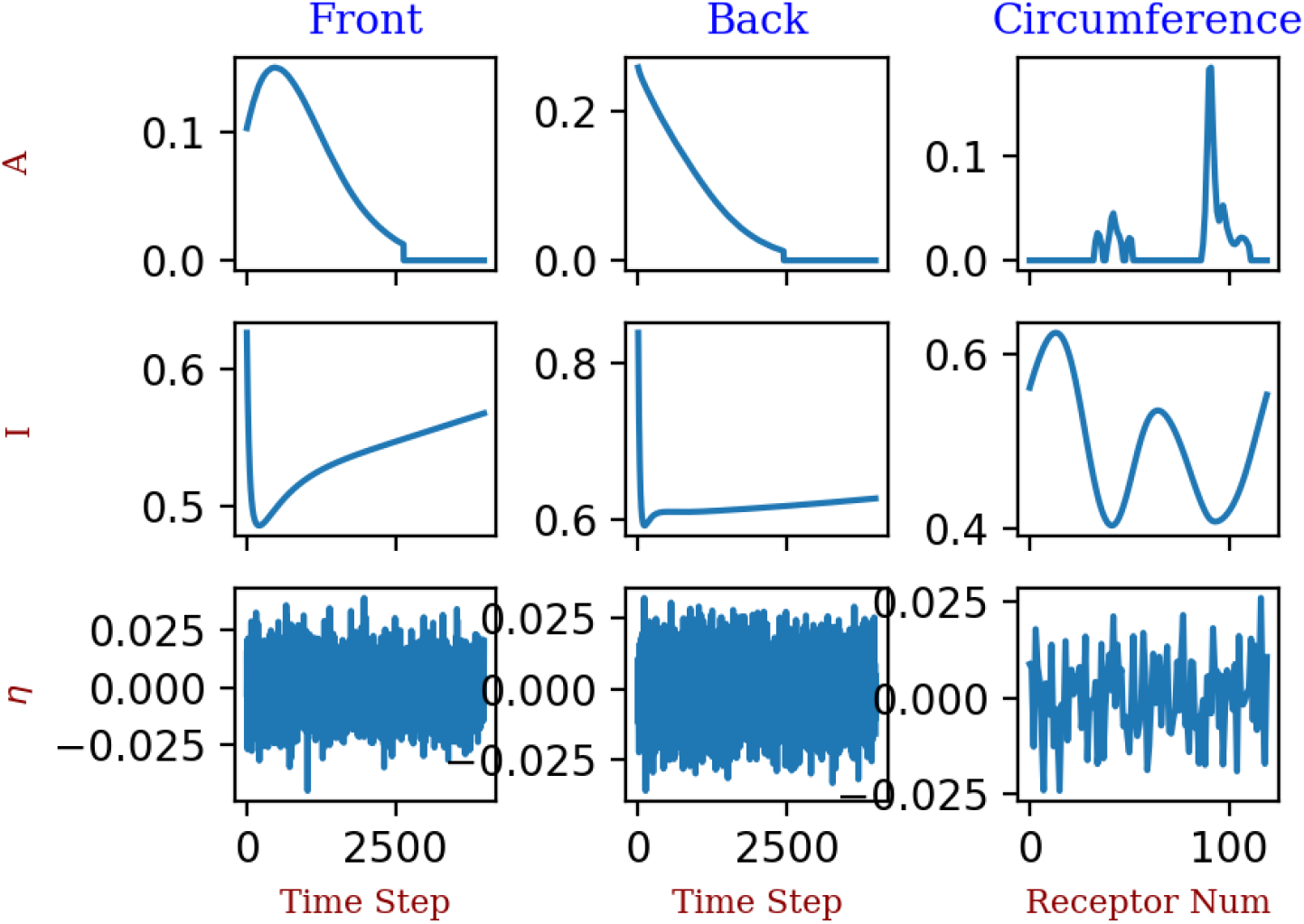
First column illustrates the concentration of *A, I*, and *η* in the “front” compartment, respectively. Second column depicts the same quantities in the “back” compartment of a cell during a single directional sensing process in a homogeneous field. Third column shows the amount of *A, I*, and *η* in all of the compartments at the almost end of a single directional sensing process, when the dominance of *A* in the compartment with index 90 is evident and its amount is going to vanish after 500 iteration. More information is in the text.

The third column of Fig.3 illustrates the corresponding concentration of *A, I*, and *η* for each compartment at the near end of the directional sensing process when the amount of *A* is going to vanish after 500 iteration. Evidently, there is a significant difference between the concentration of *A* in the “front” component and the rest of the components.

In order to find statistics of the final preferred direction by them, we perform the same simulation for 1000 cells with different initial conditions. Fig.6(a). shows the corresponding circular histogram of the final preferred direction, evaluating the frequency of repeated indices that cell front is specified by them. Naturally, we expect a uniform distribution for these statistics of a cell population located in a homogeneous medium, which seems evidently satisfied.

**Fig 4.**
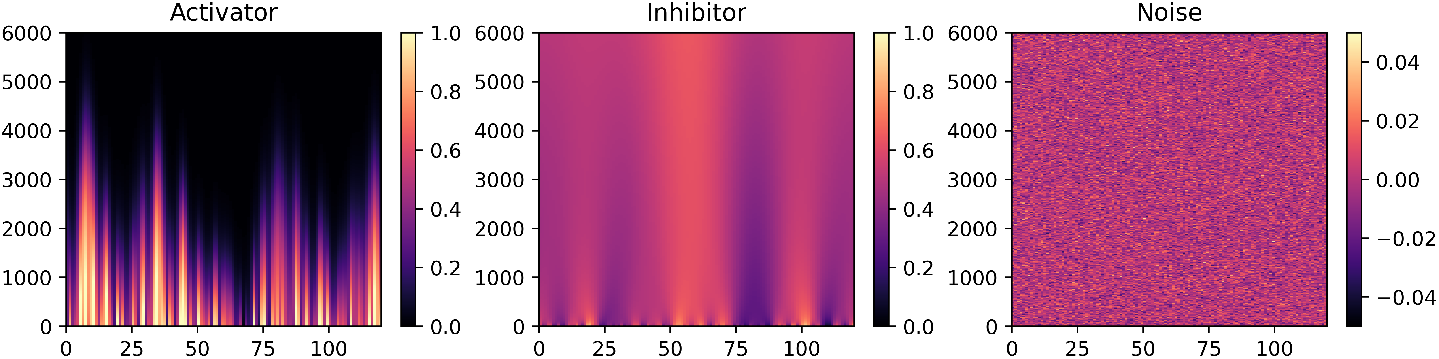
Heat map showing the concentration of (Left panel) activator concentration *A*, (Middle panel) inhibitor concentration *I*, and (Right panel) amount of noise *η* in each point of cell circumference during a single step(including 4532 iterations) in an inhomogeneous field with a right-ward horizontal linear concentration gradient (∇*c* = 0.7). Note that both index 0 and index 120 address the same compartment and point to the right direction of the horizon. Here, (a) the compartment with index 1 in which *A* outlasts more than other compartments, introduces the so-called “front” of the cell, and (b) the compartment with index 57 in which *A* outlasts more than other compartments, introduce the so-called “back” of the cell.

**Fig 5.**
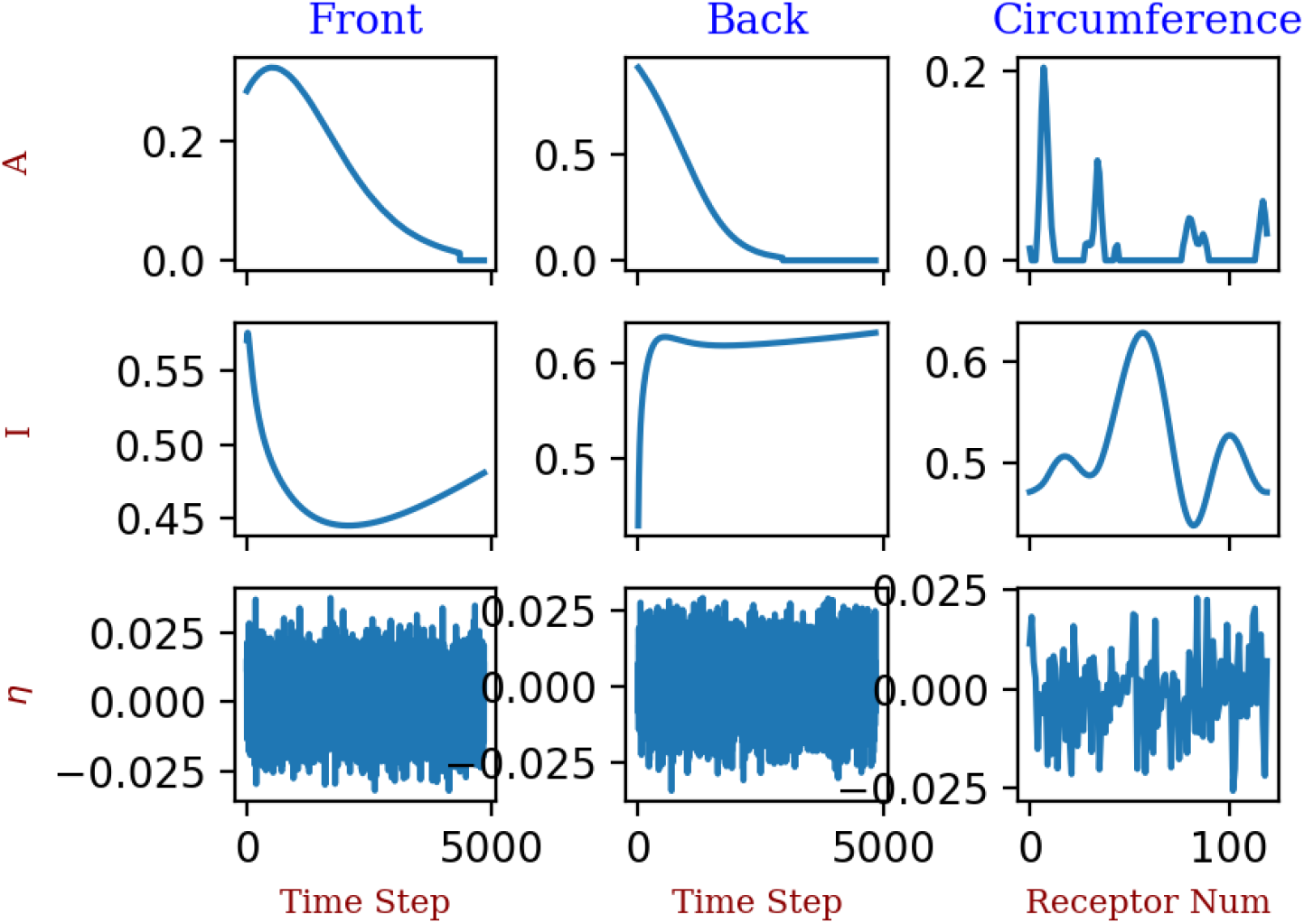
First column illustrates the concentration of *A, I*, and *η* in the”front” compartment. Second column depicts the same quantities in the “back “ compartment of a cell during a single directional sensing process in an inhomogeneous field with a right-ward horizontal linear concentration gradient (∇*c* = 0.7). Third column shows the amount of *A, I*, and *η* in all of the compartments at the almost end of a single directional sensing process, when the dominance of *A* in the compartment with index 1 is evident and its amount is going to vanish after 500 iteration. More information is in the text.

**Fig 6.**
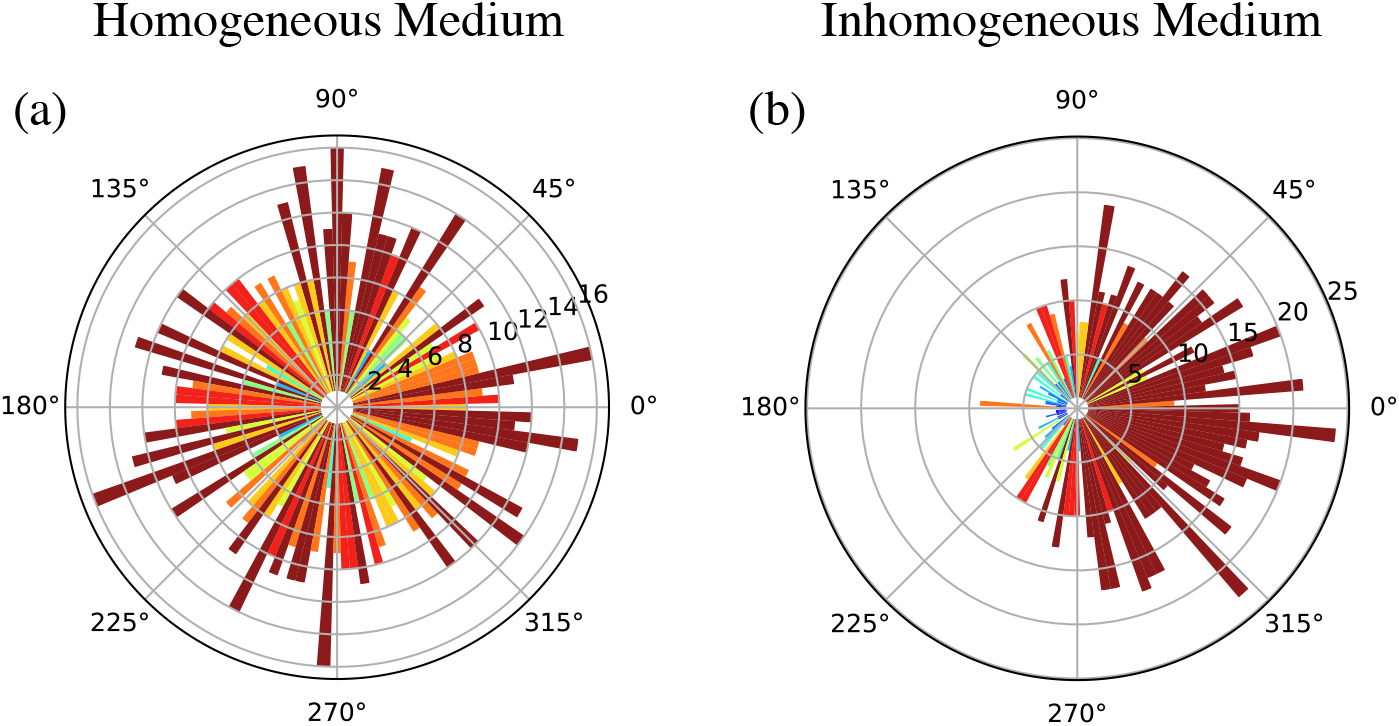
Circular histogram of preferred direction in (a) homogeneous medium in absence of chemoattractant and (b) inhomogeneous medium where ∇*c* = 0.5. The size of the statistical sample is equal to 1000

### Inhomogeneous Medium

In the presence of inhomogeneous external chemical cues, the cell’s behavior sounds similar to that of the cell in the absence of cues, except that it turns out that the highest accumulation of activators is adjusted in the direction of the concentration gradient of environmental stimulants. In the non-uniform medium, different parts of a cell are stimulated with variant concentrations of chemoattractant. This triggers downstream chemical pathways in each compartment. After sharing and comparing the local information with neighboring compartments, these complicated networks allow cells to globally detect external gradients. To study the directional sensing process in cells which are located in an inhomogeneous medium, we define an orthonormal basis with the unit vectors 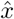 and *ŷ*, where 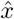 is the direction of the spatially constant gradient of chemoattractant. Moreover, we assume that the compartment with index zero is along *x* axis. Now, each compartment participates in directional sensing dynamic with what it has received from its environment at its own location. Here, we assume that the diameter of the cell is unite, without losing generality. Fig.4 depicts a typical heat map for the time evolution of activator *A*, inhibitor *I*, and stochastic variable *η* on the perimeter of a cell. Based on this simulation, the cell opts to move forward with a slight deviation (*j* = 1) with respect to *x* direction. Here, we assume that a linear concentration gradient (∇*c* = 0.7) is covered the surface on which the cell is located. The first column of Fig.5 illustrates the evolution of *A, I*, and *η* during the simulation at the front of the cell with a corresponding index of *j* = 1. The simulation takes 4532 number of iterations to give the result.The second column of Fig.5 depicts the evolution of the same variations at the rear side of the cell, with corresponding index of *j* = 57, where the amount of *I* concentration lasts more than that of other components. The third column of Fig.5 represents the concentration of *A, I*, and *η* variables at the near end of the directional sensing process of this directional sensing process. Here, the result comes out after 4032 iterations, 500 steps before the process become completed.In the presence of environmental chemical cues, cells choose their direction with lower uncertainty, and higher tenacity depending on the concentration slope of the cues, Fig.6.

## Discussion

One of the major shortfalls of a globalist view of cell, is the fact that a cell, especially of eukaryotic variety, is deeply compartmentalized, where the concentrations of different chemical species are drastically different from any two points in the cytosol.

From a hierarchical modeling insight, stimulation of this sort of cells by either environmental chemoattractants or intrinsic noise induces local downstream signaling pathways within the cell. In consequence, different segments of the cell communicate their information with each other. The interaction results in a fascinating characteristic that none of its building blocks can muster: the ability to respond to chemotactic cues present in its environment as an integrated entity. This emergent feature separates the realm of goal-less and unguided objects from the realm of living entities that change their behaviors based on cues captured from their environment. Here, we formulate the model both in the absence and presence of a gradient of environmental cues. The designed model which of course is closely linked with LEGI [14–16] and STEN [10,19,25] models, assumes a common origin for direction detection in homogeneous and heterogeneous environment. It is assumed that the signaling transduction system in a starve cell contains a network of coupled excitable compartments uniformly placed around cell circumference. Each of these component systems, consisting of two evolving variables *A* and *I*, can become excited with what it receives locally, including noise or/and signal. Neighboring compartments share information of two variables *A* and *I* with opposite effective directions: While *A* tends to become locally concentrated in a part of cell periphery, *I* is diffusing in the network. Our results show that the suggested dynamic leads to an asymmetry that lasts for only a limited period of time. This provides a potentially preferred direction for a cell to move forward along the stretch. This was less obvious from previous models, *e.g*., one of the pitfalls of incorporating *Turing* model is the fact that once the cell achieves the symmetry breaking, it is very hard to get rid of it as the asymmetric patterns are stable in this model. In order to occur patterning, the inhibitor *I* must diffuse more rapidly than the activator *A*. Thus, while activation and autocatalytic production of *A* is a localized process, inhibition of *A* by *I* is long-ranged, because once formed in the vicinity of *A, I* should rapidly diffuse away to inhibit the formation of *A* elsewhere. At the same time, this rapid diffusion of *I* ensures that it does not inhibit the local formation of *A* when it is removed from the vicinity too quickly.

On the other hand, even though it is feasible to form a dynamic pattern with the help of excitable systems, in order to have spatiotemporal patterns, it is necessary for activator variable to move faster than the inhibitor one [26],which is not the case in reality.

In our model, we incorporated the main features of excitable systems of Barkley type [22], except that here not only the activators do not spread out, but also some sort of piling up in a localized spot, near which concentration of stimulants are higher, is favored by them. In order to make it come true, one can presume that the activator variable *A* has something to do with the actin polymerization process. One can envisage that the chemical equilibrium of Actin polymerization ⇋ Actin Depolymerization shifts to the left side on the spots where the amounts of chemoattractants are higher. As a consequence, actin monomers, which are present everywhere in the cytosol, set out for the points where polymerization is more likely. It has been shown that there are biochemical agents, *e.g*., *PH* domain proteins, that translocate to the front of cell during the formation of the protrusions to promote actin polymerization at that area. Likewise, there are biochemical agent,e.g., *PTEN*, that inhibit pseudopod formation at the sides and interacts with myosin II contraction at the rear side of cell [27]. Although in our model *A* and *I* do not stand for real biochemical components, this approach lets us explore the logic of directional sensing phenomenon better and deeper. Indeed, having this sort of dynamics in mind, one may construct a LEGI model on top of them which satisfies most of the requirements of a proper directional sensing model, including adaptation as well. Evidently, the external concentration gradient of chemoattractants directs the system to have more protrusions on the front than in the back [28, 29]. The compartmentalized model simply exhibits this as the gradient would displace the steady state such that becomes easier to excite in front and less easy in the back. Fig.6 illustrates the statistics of the preferred direction of independent realizations of the proposed dynamic for directional sensing both in the absence (a) and presence (b) of chemoattractants. The statistics share a high resemblance to that of the chemotactic index, a measure of the average angle of propagation relative to the external gradient direction of the stimulants, of a motile cell. Except that, in Fig.6 all the realizations are independent, meaning that the initial values of activator variable *A* and inhibitor variable *I* are randomly picked up from the interval (0,0.5). However, in the calculation of the chemotactic index there are correlations between successive steps in a trajectory along which an individual cell moves on,i.e. new pseudopodia frequently emerge while the previous pseudopodium has not vanished yet [30]. This implies some sort of memory in directional displacement as the initial values of the activator and inhibitor of consequent steps are not independent. In the case of a homogeneous medium, it has been shown that the subsequent direction depends not only on the present direction, as in a standard Markov process, but also on the previous direction [29]. However, the short memory of the second order Markov chain vanishes on the cell’s stationary state and leads to diffusive behavior for squared displacement.

Our model, like every excitable model, does also a good job in the amplification of external signals. This is because the crimson-gray excursion in Fig.1(d) is long enough to intensify the response and thus the system would have a large amplification. Moreover, in the model, we benefit from a switch-like threshold *δ* between excited and resting states. This algorithm not only provides both simplicity and efficient numerical implementation [22], but also suggests a methodical way for including activation-bias and adaptation capability in an excitable system based on spatial heterogeneities [31]. In this model, we took advantage of a “mean-reverting” process to provide the included noise in the system dynamic [24]. The noise is of a colored type, arising from an Ornstein-Uhlenbeck process. The process is a Markov one, since it includes an uncorrelated Gaussian noise and, thus, does not depend on the history of the process. The correlation time of the process decays exponentially, see Fig.1(e) inline. The *λ* parameter has time inverse dimension and indicates the cut-off frequency of the noise fluctuations. Similar dynamics in a two-dimensional system was used to model orchestrated signaling the cells [32]. Recently, it has been shown that amoeboid and fan-shaped modes can be seen as coexisting behavioral traits of the same cell rather than signatures of heterogeneity in a cell population, where different modes of motility occur in different cells due to cell-to-cell variability [33]. In our future work, we are about to go through the parameter space of the offered model in this manuscript and assay the parameters upon which the coexisting as well as other possible behaviors of cells may turn out.

## Code and Data availability

The software used to run all simulations was Python. The scripts are available from the corresponding author upon request.

## Author contribution

M.S and Z.E implemented research concept and designed modeling framework. Z.E performed numerical simulations and interpreted the results. Z.E wrote original version the manuscript. Z.E and M.S contributed to manuscript revision.

## Additional information

The authors declare no competing interests.

